# Gene regulatory network inference from perturbed time-series expression data via ordered dynamical expansion of non-steady state actors

**DOI:** 10.1101/007906

**Authors:** Mahdi Zamanighomi, Mostafa Zamanian, Michael Kimber, Zhengdao Wang

## Abstract

The reconstruction of gene regulatory networks from gene expression data has been the subject of intense research activity over the past many decades. A variety of models and methods have been developed to address different aspects of this important problem. However, these techniques are often difficult to scale, are narrowly focused on particular biological and experimental platforms, and require experimental data that are typically unavailable and difficult to ascertain. The more recent availability of higher-throughput sequencing platforms, combined with more precise modes of genetic perturbation, presents an opportunity to formulate more robust and comprehensive approaches to gene network inference. Here, we propose a step-wise framework for identifying gene-gene regulatory interactions that expand from a known point of genetic or chemical perturbation using time series gene expression data. This novel approach sequentially identifies non-steady state genes post-perturbation and incorporates them into a growing series of low-complexity optimization problems. The governing ordinary differential equations of this model are rooted in the biophysics of stochastic molecular events that underlie gene regulation, delineating roles for both protein and RNA-mediated gene regulation. We show the successful application of our core algorithms for network inference using simulated and real datasets.

## I. Introduction

The elucidation of gene regulatory networks is fundamental to understanding the dynamic functions of genes in biochemical, cellular and physiological contexts. The architectures of networks comprised of small numbers of genes are generally deciphered using classical experimental techniques, where biophysical data describing the interactions of genes and their products can lead to useful models and well-characterized systems. While this validated experimental tract continues to provide valuable biological insight, it is ultimately laborious and costly, and often demands strategies uniquely tailored to individual biological systems and problems. Furthermore, the models that result from these efforts tend to be limited to a very modest subset of genes, typically suffer from a lack of temporal resolution, and focus narrowly on very particular modes of interaction.

To complement these established approaches, there is a great impetus to develop more efficient and uniformly applicable *in silico* methods for gene network inference and discovery [1], [2], [3], [4], [5], [6], [7]. Of particular interest is the goal of gene network inference using perturbed gene expression data [8], [9], [10], [11], [12], [13], [14], [15], [16] whereby gene expression levels are measured under the influence of either genetic or chemical perturbations of the system. Previous attempts at network reconstruction via perturbation tend to be limited to the analysis of steady-state gene expression. The growing ubiquity of next-generation sequencing technologies presents a powerful high-throughput substrate for capturing the dynamic and non steady-state aspects of gene expression.

In this work, we seek to develop a robust framework for network inference that relies on temporal gene expression data coupled to genetic or chemical perturbation. In a departure from previous attempts, our formulation does not require *a priori* knowledge beyond the set of temporal gene expression measurements, acknowledges the non-steady state and dynamic nature of gene expression, incorporates both RNA and protein-mediated regulation, sequentially absorbs a growing number of genes into the regulatory network immediate to perturbation, aims for sparsity in network topology, and reduces an otherwise complex optimization problem into a convex form that can be solved efficiently.

### Notation

Throughout this paper {*d, i, j, k, l*} count integer numbers. Column vectors and matrices are indicated by bold lower-case and upper-case letters, respectively. We use 1 to show a vector with all entries 1 and 0 a vector with all entries 0. The set of real numbers is denoted as ℝ and positive real numbers ℝ^+^. The indicator function 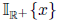 has the value one when *x* ∊ ℝ^+^, otherwise zero. The operator sign(**x**) replaces each entry of **x** with its sign function value. We use (**X**)^*T*^ to denote transpose of **X**, *dx*(*t*)/*dt* and *x*′(*t*) the first derivative of *x*(*t*) with respect to time *t*, ||**x**||_1_ the 1-norm of vector **x**, ||**x**||_2_ the 2-norm of vector **x**, and ||**X**|| the largest singular value of matrix **X**. We explicitly state a function of time in the form **x**(*t*). This is to be distinguished from vectors of the form **x**(*i*), where *i* is a positive integer representing the *i*th entry of the vector **x**.

## II. System Model

### A. *Gene expression datasets and perturbation*

Let *x*_*i*_(*t*) and *y*_*i*_(*t*) denote the RNA-level and protein-level expression of gene *i* at time *t*, respectively. We define an *m* × *n* gene expression matrix

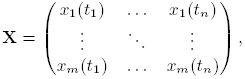

where *m* indicates the total number of genes in the system and *n* the total number of samples in the time series. In practical cases, with expression data originating from microarray or RNA-Seq experiments, *m* ≫ *n*.

The paper is concerned with datasets with known points of perturbation. In this experimental scheme, a gene 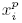 is specifically targeted for perturbation via either gene suppression or gene overexpression. Perturbation is triggered at a known time point after a series of presumably steady state measurements. Without loss of generality, it is assumed that the starting point of perturbation occurs at *t*_1_ and prior measurements are approximately steady state. Datasets from experiments that conform to this scheme are in the following form, where 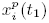 represents the point of perturbation and *L* denotes the total number of samples post-perturbation.

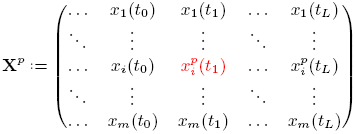

### B. *Conceptual description of inference approach*

We consider a non-perturbed system as one with genes in steady state, i.e., where *dx*_*i*_(*t*)/*dt* and *dy*_*i*_(*t*)/*dt* are approximately zero. After a series of steady state expression measurements, a protein-encoding gene in this system is perturbed to bring about a dramatic change in its expression level, i.e., where 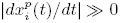, followed by a series of post-perturbation measurements. The discrete set of expression measurements, with appropriate temporal resolution, can be used to produce continuous gene trajectory curves.

For a short period of time post-perturbation, the perturbed gene falls out of steady state while all other genes remain effectively in steady state. The induced change in RNA expression, 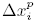, is coupled to a delayed change in protein expression, 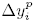. This shift in protein availability leads, through the immediate regulatory network of the perturbed protein, to changes in the expression levels of other genes.

Consider the set of all genes that are affected by 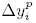 at time *t*. We divide this set into protein and miRNA-encoding subsets. The set of all indices that correspond to protein-encoding genes is shown as as *G*(*t*), and *M*(*t*) is set of all indices that correspond to miRNA-encoding genes. We define the collection of RNA expression data for these subsets as *χ*_*G*(*t*)_ := {*x*_*i*_(*t*)|*i* ∈ *G*(*t*)} and *χ*_*M*(*t*)_ := {*x*_*i*_(*t*)|*i* ∈ *M*(*t*)} respectively. We further define the collection of protein expression levels for subset *G* as *Y*_*G*(*t*)_ := {*y*_*i*_(*t*)|*i* ∈ *G*(*t*)}.

In principle, we can identify genes that fall out of steady state in an ordered manner with gene trajectory analysis. The growing set of non-steady state actors in the system, both members of *G*(*t*) and *M*(*t*), can then be sequentially incorporated into a growing network of interactions to be modeled.

### C. *Governing regulatory equations*

Gene and protein expression dynamics are often modeled in the form of ordinary differential equations [17], [18], [19], with gene-specific rate constants for molecular synthesis and degradation and gene-specific functions accounting for the regulatory effects of proteins. We introduce miRNA-mediated gene regulation into this model and establish functions for both protein and RNA regulatory interactions that complement our overall approach to network inference. The architecture of the gene regulatory circuit under consideration is depicted in Figure 1.

**Fig. 1.**
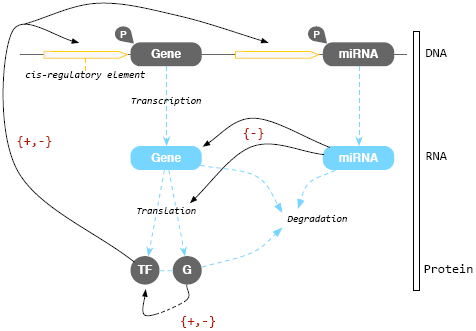
Gene regulatory circuit. ‘Gene’ represents protein-encoding genes and ‘miRNA’ represents miRNA-encoding genes. Protein-encoding genes can give rise to transcription factors (‘TF’) that directly exert influence on the cis regions of other genes, as well as non-TF proteins (‘G’) that can indirectly act through TFs and various biochemical cascades. These protein regulators ultimately affect the equilibrium probability of RNA polymerase (‘P’) being bound to a promoter of interest. Additionally, miRNAs can directly repress expression via targetted RNA degradation or translational repression. All proteins and RNAs in this system undergo varying rates of chemical degradation.

This circuit can be represented in the following form:

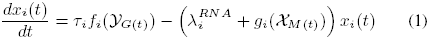

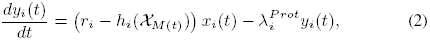

where *τ*_*i*_ is the rate of transcription when RNA polymerase (RNAP) is bound, *f*_*i*_(*y*_*G*(*t*)_) is the probability of RNAP binding, 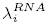 is the rate of basal RNA degradation, *g*_*i*_(*χ*_*M*(*t*)_) incorporates the effect of miRNA-mediated RNA degradation, *r*_*i*_ is the rate of translation, *h*_*i*_(*χ*_*M*(*t*)_) accounts for the effect of miRNA-mediated translational inhibition, and 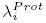 is the rate of protein degradation. It follows from the biological definitions of the system that parameters *τ*_*i*_, 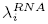, *r*_*i*_, and 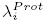 are to be positive and *h*_*i*_(*χ*_*M*(*t*)_) ≤ *r*_*i*_.

### D. *Protein-mediated regulation*

For each gene, *i*, we employ an existing statistical thermodynamic framework [20], [21] to model the equilibrium probability of RNAP binding to a gene of interest as a function of protein regulators, *f*_*i*_(*y*_*G*(*t*)_). We extend a previous derivation of multiple protein regulators operating on a single gene [22] and explicitly show that the general form can be expressed as a function of non-steady state genes, *G*(*t*) (Appendix A). Although steady state regulators play an active role in gene regulation, we can effectively restrict our binding probability function to the activities of perturbed regulators. This function is shown below.

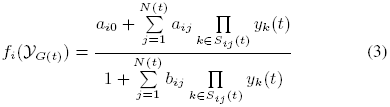

where *S*_ij_(*t*), 0 ≤ *j* ≤ *N*(*t*), is the list of all possible protein products of genes within set *G*(*t*) that interact to form regulatory complexes. For instance when *G*(*t*) = {1, 2}, there are *N*(*t*)+1=4 complexes as the empty set *S*_*i*0_ = {∅}, *S*_*i*1_ = {1}, *S*_*i*2_ = {2}, and *S*_*i*3_ = {1,2}. To reduce the complexity of this model, we restrict *S*_*ij*_ (*t*) to all terms up to the second-order, accounting for the interactions of no more than two proteins bound together. In this arrangement, a complex represents either the products of a single gene or the interaction of the products of any two genes that can form a regulatory agent. However, any number of complexes can additively combine to regulate single genes. The numbering of complexes is an arbitrary labeling of genes and gene-pairs in the system. The coefficients 0 ≤ *a*_*ij*_ ≤ *b*_*ij*_ depend on the binding energies of regulator complexes that act on a promoter region, and *a*_*i*0_ and *b*_*i*0_ correspond to the case where no regulators are bound to the promoter region (∏_*κ∈sio(t)*_*yk(t)* :=1). It is assumed all coefficients are normalized so that *b*_*i*0_ = 1.

### E. *E. miRNA-mediated regulation*

To account for the effects of miRNA regulation on each gene, we draw on previous mass-law (linear) models [23], [24] that acknowledge two primary routes of inhibitory regulation: (i) cleavage or degradation of target transcript and (ii) translational repression. These are represented by functions *g*_*i*_(*χ*_*M(t)*_) and *h*_*i*_(*χ_M(t)_*), respectively. The former is a modifier of the RNA degradation rate constant, 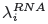, while the latter detracts from RNA available to the translational machinery without affecting RNA concentration as assayed. These functions are shown below.

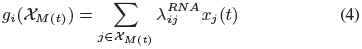

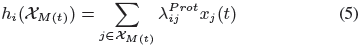

where both 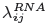 and 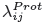 are greater than or equal to zero.

We impose the constraint that any given miRNA can only inhibit the expression of a particular target mRNA through one mode of regulation, either transcript cleavage or translational repression This is reasonable, given that the particular pathway of inhibition is determined by the specificity of binding between a particular miRNA and a seed site on a target transcript, which is a fixed interaction for each miRNA-mRNA pairing [25], [26], [27]. This constraint takes the following mathematical form

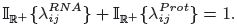

## III. Network Inference Algorithm

Sub-sections III-A - III-D contain all the core algorithmic components in our proposed inference pipeline. A graphical overview of how these modular algorithms form a framework for gene network inference is shown in Figure 2. This linear ordering of post-processing and inference steps, although designed for a normalized gene expression dataset involving a precise perturbation, is robust and flexible.

**Fig. 2.**
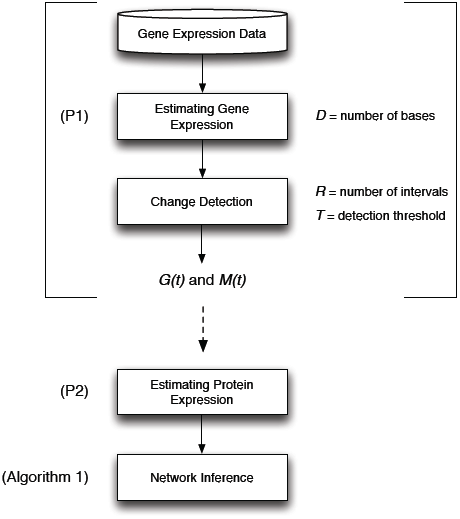
Overview of gene inference pipeline, beginning with a normalized gene expression dataset. The first stage involves the estimation of all gene trajectories as noise-free and continuous curves (P1), followed by segmentation into equally-spaced intervals for detection of significant changes in expression. The time-dependent expansion of *G*(*t*) and *M*(*t*), along with the result of (P1), seed downstream network inference. In the next stage, (P2) is used to estimate protein expression, and finally all obtained results are considered in algorithm 1 to produce a regulatory network map. Figure 3 provides a graphical description of the bracketed pre-inference stages.

**Fig. 3.**
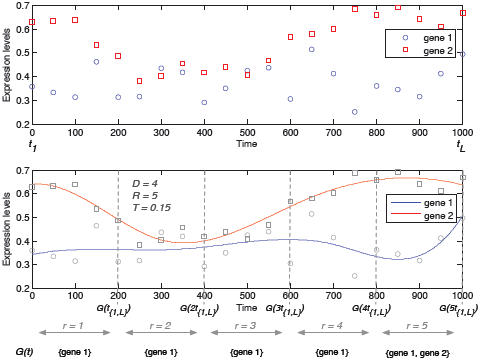
The bracketed pre-inferenced stages of the pipeline in Figure 2 are shown graphically. Discrete expression data from two genes and a small number of basis functions are utilized to produce continuous models of expression (P1), followed by segmentation and change detection. In this simple example, a change in gene 2 is detected in sub-interval *r* = 1, and a change in gene 1 is detected in sub-interval *r* = 5.

### A. Modeling ami estimation of gene expression

Normalized gene expression values, such that *x*_*i*_(*t*) ≤ 1, are the given input for the algorithms described in this and subsequent sections . In reality, gene expression trajectories are inevitably noisy, which perturb the model parameters away from the true values. To reduce this noise effect, we first represent gene expressions as a linear combination of basis functions in the following form

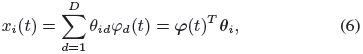

where *D* is the total number of bases and *θ_id_* the coefficient of the dth basis function, *φ_id_*(*t*). The basis functions are chosen to take the form of a B-spline (Appendix B). Although all genes are associated with a common set of basis functions in (6), one can consider different sets of basis functions for different genes.

The form of (6) allows us to fit a continuous function for a set of discrete gene expression measurements, using the following minimization

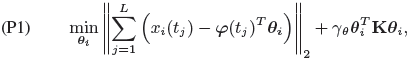

where the roughness penalty 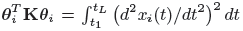 and **K** is a roughness matrix with the (*j,k*)th entry 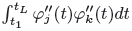. Here, the first term is intended to diminish noise within measurements and the second term is intended to smooth our approximations. The parameter *γ_θ_* is tuned by cross validation where training data is available, otherwise it can be drawn from a characterized network from the nearest available biological system.

Employing (P1), our estimation to *x*_*i*_(*t*), denoted as 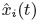, is a continuous function in time and its first derivative can be easily calculated as

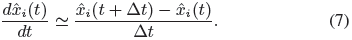

Throughout the rest of the paper, it is assumed that our samples are taken from 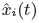 and therefore, any arbitrary number of samples, *L*, is achievable. We further replace 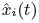 with *x*_*i*_(*t*) for notational convenience.

### B. Detection of perturbed genes

We can introduce a simple first approach for detecting when individual genes exit steady state post-perturbation. Gene expression models generated via (P1) are essentially smooth and noise-free when the total number of bases is restricted to an appropriately small number, *D*. High-frequency gene trajectories, whether a product of noise or periodicity in expression [28], [29], are converted into flat trajectories. This property allows us to detect when significant non-periodic deviations occur with respect to the initial steady state measurement(s). More precisely, time interval [*t*1, *t*_*L*_) is divided into *R* sub-intervals as [*rt*_{1,*L*}_, (*r* + 1)*t*_{1,*L*}_) for all 1 ≤ *r* ≤ *R*, where *t*_{1,*L*}_ := (*t*_1_ — *t*_*L*_)/(*R*+1). We choose *R* with respect to the nature of the original expression data, such that *R* ≥ *D*.

For each sub-interval, we look for the maximum and minimum values of trajectories. The sets *G*(*t*) and *M*(*t*) are then expanded as follows. At sub-interval *r*, gene *i* is included within either *G*(*t*) or *M*(*t*) for *t* > *rt*_{1,*L*}_ provided that the deviation from the steady state measurement of gene *i* is greater than a desired threshold, *T*. In the simulations described in this paper, *T* was set in the range of [0.15, 0.20] for normalized expression data. Both *R* and this threshold can be modified to better reflect the frequency of gene expression measurements for a given biological system. If more complex change detection schemes are preferred, a number of alternative approaches can be adapted for this purpose [30], [31], [32].

### C. Modeling and estimation of protein expression

**Formulation:** Similar to (6), we express the protein level *y*_*i*_(*t*) as

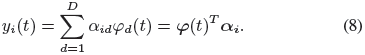

Our objective is first to find *α_i_* through the ODE (2) resulting in an estimation of the protein level *y*_*i*_(*t*). The calculated *y*_*i*_(*t*)’s are in turn used to approximate unknown variables associated with the ODE (1). One of the challenges of solving non-linear ODEs is that the solution does not usually have a closed form. We propose to transform the non-linear ODE (2) into a linear regression problem. To motivate our method of constructing the ODE solution, we consider the first derivative of *y*_*i*_(*t*) as

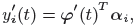

and ODE (2) is consequently represented as

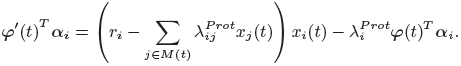

We rewrite the above equation in the following form

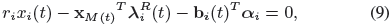

where 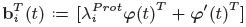 and 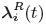 is the column vector with entries 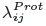, ∀*j* ∈ *M*(*t*). The miRNA expressions corresponding to 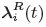 are indicated by the vector **x**_*M*(*t*)_ such that both vectors, 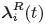 and **x**_*M*(*t*)_, have the same index order. For notational convenience, we assume that all entries of **x**_*M*(*t*)_ are multiplied by *x*_*i*_(*t*).

Consider gene expressions at times *t*_*l*_, 1 ≤ *l* ≤ *L*. Setting all available gene expressions in equation (9), we arrive at

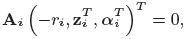

where

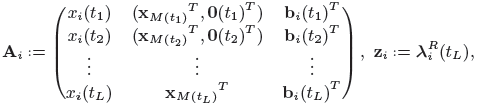

and **0**(*t*_*l*_) is the zero column vector with length **card**(*M*(*t*_*L*_)) — **card**(*M*(*t*_*l*_)). When the length is zero, we do not consider the vector **0**(*t*_*l*_), e.g., 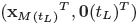 is replaced by 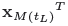 in the last row of **A**_*i*_. Matrix **A**_*i*_ has *L* rows and **card**(*M*(*t*_*L*_)) + *D* + 1 columns. Given that *r*_*i*_ is positive, we normalize 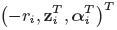 with respect to *r*_*i*_ and represent the normalized vector as 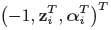, acknowledging abuse of notation. Given 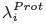 and *M*(*t*), matrix **A**_*i*_ is completely determined.

**Algorithm:** We need to solve the linear system model

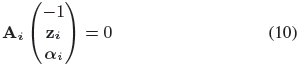

for **z**_*i*_ and **α**_*i*_ when matrix **A**_*i*_ is determined. For identifiability of **z**_*i*_ and **α**_*i*_, we require that *L* ≥ **card**(*M*(*t*_*L*_)) + *D*, that is the number of equations is no smaller than the number of unknown parameters. However the sparsity in **z**_*i*_, given that only a small number of miRNAs typically act on a common gene [33], reduces the number of required equations.

To account for measurement noise and encourage **z**_*i*_ to be sparse, we will minimize the 2-norm error described in (10) with 1-norm regularization ∥z_*i*_∥_1_. Furthermore, we adopt the analogous roughness penalty 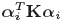 as used in (P1). Thus, we propose to obtain the ODE (2) solution with the following convex optimization

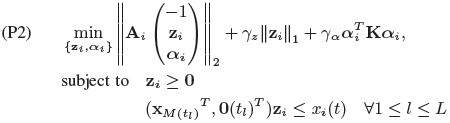

where *γ_z_* and *γ_α_* are chosen using cross validation. The second constraint ensures that the total rate of translation, *r*_*i*_ — *h*_*i*_(*χ*_*M*(*t*)_), is not negative. Due to the convex nature of this problem, it can be quickly solved for large gene datasets. This recovery of protein expression is dependent on prior knowledge of individual protein degradation rates, 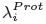. In the absence of this experimental data, we can fix the value of 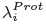 to 1 for the entire system and still achieve accurate network reconstruction as shown in subsequent sections.

### D. Gene regulatory inference

**Formulation:** The model given by ODEs (1) and (2) describes the evolution of RNA and protein expressions provided that we know all the regulatory parameters, e.g., *a*_*ij*_, *b*_*ij*_, and *τ_i_*. Coefficients *a*_*ij*_ and *b*_*ij*_ are difficult to experimentally determine and it is currently infeasible to carry out the relevant measurements simultaneously for a complex system with a large number of genes and gene products under consideration. Our goal is to estimate these coefficients so that the ODE models can be temporally fitted to large gene expression data. Specifically, we will use the previously described estimations of protein and RNA expression to approximate *a*_*ij*_ and *b*_*ij*_, and to infer a regulatory network map.

To improve the reliability of the inferred network, we take into account time-dependent changes in gene levels and construct a set of equations accordingly. This is an important departure from standard steady state treatments. In this scenario, we first assume that the non-perturbed system is in an initial steady state, where RNA and protein levels are near constant (i.e., *dx*_*i*_(*t*)/*dt* = *dy*_*i*_(*t*)/*dt* ≃ 0). As previously mentioned, the perturbation of protein-encoding gene 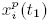 first leads to fluctuations in the expression levels of genes in its immediate regulatory network. Genes that have exited a steady-state expression profile at any time up to *t*, *G*(*t*) and *M*(*t*), expand to contain greater numbers of genes that interact to form a putative regulatory network.

Considering changes in gene levels *x*_*i*_(*t*) at time *t*_*l*_, 1 ≤ *l* ≤ *L*, with the exception of 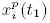, the term τ_*i*_*f*_*i*_(*y*_*G*_(*t*_*l*_)) in equation (1) can be rewritten as follows

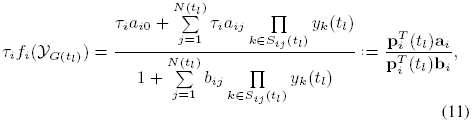

where **α**_*i*_ is a vector with (*j* + 1)th entry *τ_i_a_ij_*, 0 ≤ *j* ≤ *N*(*t*_*L*_). The (*j* + 1)th element of vector **p**_i_(*t*_*l*_) is described by ∏_*κ∈s_ij_(t*_*l*_)_*yk(t)* when 0 ≤ *j* ≤ *N*(*t*_*l*_) and zero for *N*(*t*_*l*_)+1 ≤ *j* ≤ *N*(*t*_*L*_). Vector **b**_i_ is defined such that the first entry is *1* and (*j*+1)th, 1 ≤ *j* ≤ *N*(*t*_*L*_), is *b*_*ij*_.

#### Remark 1.

*Given that y_i_*(*t*)*s are normalized with respect to r_i_, a_ij_ and b_ij_ include the multiplier term ∏_*κ∈sio(tl)*_^*rk*^ so that the normalization can be vanished. Similarly, τ_i_ can be absorbed into the coefficients a_ij_, where we assume τ_i_ < 1 to maintain the algorithm constraint 0 ≤ a_i_ ≤ b_i_.*

We also represent

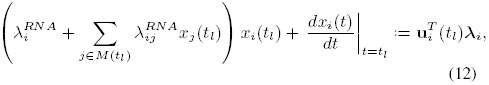

in which **u**_*i*_(*t*_*l*_) and **λ**_*i*_ are defined as follows. First and second entries of vector **u**_*i*_(*t*_*l*_) are *dx*_*i*_(*t*)/*dt*/_*t=tl*_ and *x*_*i*_(*t*_*l*_), respectively. The remaining entries are *x*_*j*_(*t*_*l*_)*x*_*i*_(*t*_*l*_), *j* ∈ *M*(*t*_*l*_). Making the same arrangement of array as **u**_*i*_(*t*_*l*_), vector λ_*i*_ is determined by first entry 1, second entry 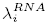, and subsequent entries 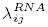, *j* ∈ *M*(*t*_*l*_).

Using (11)-(12), equation (1) can be reformulated as

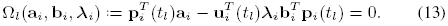

**Algorithm**: We need to solve the non-convex problem

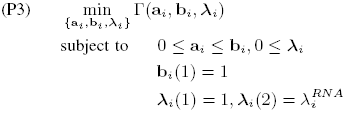

with

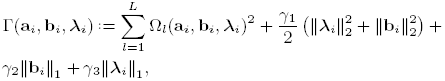

The first term in the above equation follows from (13). The second term associated with *γ*_1_/2 motivates grouping effect among variables **b**_*i*_ and **λ**_*i*_ [34], [35]. Due to the assumption that each gene has only a few regulators, 1-norm regularizations are considered to encourage sparse solutions. Note that in the absence of miRNAs (all 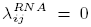, terms ∥*λ*_2_∥ and ∥*λ*_1_∥ are no longer needed.

Non-convex optimizations are generally hard to solve in a reasonable time. Hence, we seek to identify a special treatment that reduces the computational complexity and provides desired solutions. Optimization (P3) is convex in {**a**_*i*_, **b**_*i*_} for fixed **λ**_*i*_ and vice versa, and therefore the problem is bi-convex and can be solved using a variation of the alternating-direction method of multipliers (ADMM) which cycles over two groups of variables [36], cf. Appendix C. Here, given the absence of dual variables, ADMM is reduced to simple alternating minimization. The proposed solver entails an iterative procedure compromising two steps per iteration *k* = 1, 2,…

#### **Algorithm 1** : Gene regulatory inference

**input a**_*i*_, **b**_*i*_, **λ**_*i*_

**initialize a**_*i*_[0], **b**_*i*_[0], and **λ**_*i*_[0] at random with respect to

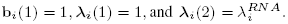

**for** *k* = 0,1,… **do**

**[S1] Update primal variables a**_*i*_ **and b**_*i*_:

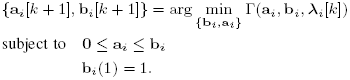

**[S2] Update primal variable λ*i*:**

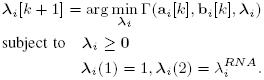

**end for**

**return a**_*i*_, **b**_*i*_, **λ**_*i*_

This iterative procedure implements a block coordinate descent method [37]. At each minimization, the variables that are not being updated are treated as fixed and are replaced with their most updated values. Then the iteration alternates between two sets of variables, {**b**_*i*_, **a**_*i*_} and **λ**_*i*_.

One difficulty with the proposed solver is that it may result in stationary points which are not necessarily globally optimal. This occurs since optimization (P3) is not convex in {**b**_*i*_, **a**_*i*_, **λ**_*i*_}. Motivated by the proposition 1 in [38], the next theorem offers a global optimality certificate upon the convergence of the solver.

#### Theorem 1.

*Let* 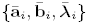 *be a stationary point of (P3). If*

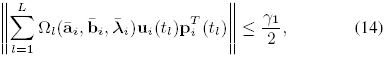

*then 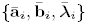 is the globally optimal solution of (P3).*

*Proof*. See Appendix D.

#### Remark 2.

*For non-convex problems, ADMM offers no convergence guarantees. Nevertheless, there are evidences in the literature that show empirical convergence of ADMM, particularly when the nonconvex exhibits specific structures. For example in our scenario, problem (P3) is bi-convex and admits unique closed form solutions for sub-problems **[S1]** and **[S2]**. This observation along with desired properties, Theorem 4.5 and 4.9 in [39], are indeed a sufficient case for successful convergence. A formal proof of convergence is beyond the scope of this paper.*

Algorithm 1 is intended for the case in which the RNA degradation rates, 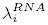, are available. However, experimentally measuring 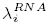 is a difficult task. We offer a simple modification to the algorithm so that network inference can be still obtained without prior knowledge of RNA degradation rates.

For simplicity of explanation, we can first remove miRNAs from our model. ODE (1) can then be rewritten as

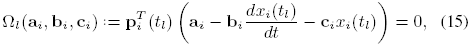

and 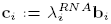. Employing the above reformulation, unknown variables **a**_*i*_, **b**_*i*_, and **c**_*i*_ are estimated through the following convex optimization

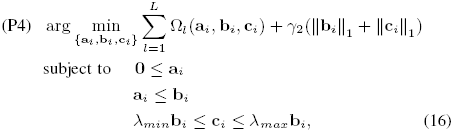

where *λ_min_* and *λ_max_* specify an lower and upper bound for 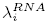, respectively. Variable **c**_*i*_ is introduced to remove 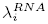 from our optimization. However, the new variable expands the feasible set of solutions, which might create an answer different from the true value. To reduce this effect, we add constraint (16) to (P4) to tighten the feasible set of solutions. Given that 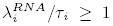, we can take on the additional constraint **a**_*i*_ ≤ **c**_*i*_. In the subsequent simulations, **λ**_*min*_ is in the near-zero range [0.001, 0.01], and **λ**_*max*_ is selected in the range [0.1,1]. It is straightforward to generalize the introduced approach within the framework of (P3). Derivations are removed to avoid repetition in the paper.

## IV. Simulations

### A. Small gene network with prior knowledge of degradation rates

To demonstrate the proposed time-series approach, we consider the three-gene network described by the following systems of ODEs for gene expression

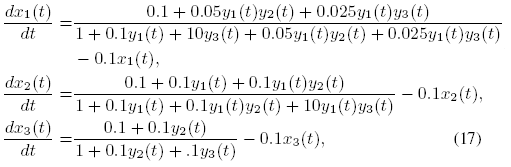

and the following system of ODEs for protein expression

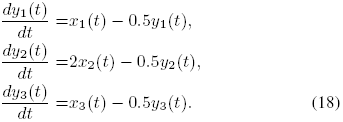

The above toy model, visualized in Figure 4, is provided to better explain our algorithms. Although a small network is examined, many of the same qualitative characteristics of large network are investigated in this example. The explicit system of ODEs, describing the kinetics of the system [40], allows us to generate samples to fit our model and to also compare recovered solutions with the ground truth. This model also incorporates complex modes of regulation, including self-regulation and combined regulators.

**Fig. 4.**
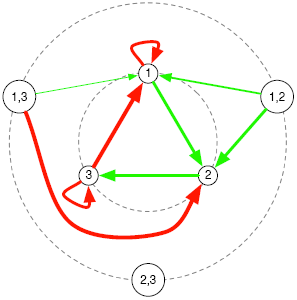
Map of gene regulatory network described by equations (17) and (18). First-order (single) and and second-order (combined) regulators are depicted in concentric circles. Green arrows specify gene activation and red arrows specify gene repression. The relative magnitudes of activation and repression are roughly represented by arrow thickness.

To generate data, arbitrary initial conditions are assigned to ODEs (17) and (18) and the system is allowed to resolve to a steady state. To perturb this steady state, the expression level of gene 1, *x*_1_ (*t*), is artificially fixed to 0.3, leading to fluctuations in the expression levels of other genes. Figure 5 illustrates expression trajectories before and during the perturbation

**Fig. 5.**
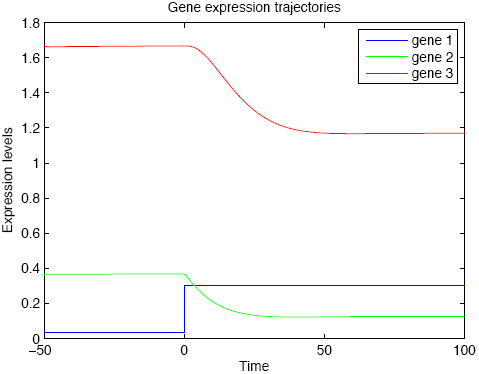
Gene expression trajectories (unnormalized) before and during the imposed perturbation. The system is in steady state before time 0. Gene 1 is artificially perturbed at time zero, leading to changes in gene expression levels. A new steady state is eventually achieved at approximately time 50. We sample expression levels between time 0 (the starting point of perturbation) and 50 (the new steady state) and use them as data in our algorithm.

We collect 12 samples from each gene expression level. The samples are chosen uniformly from time interval [0, 50]. Points 0 and 50 specify the times at which the perturbation starts and the system reaches a new steady state, respectively. using these sampled data, we solve optimization (P2) to effectively recover protein expressions as shown in Figure 6

**Fig. 6.**
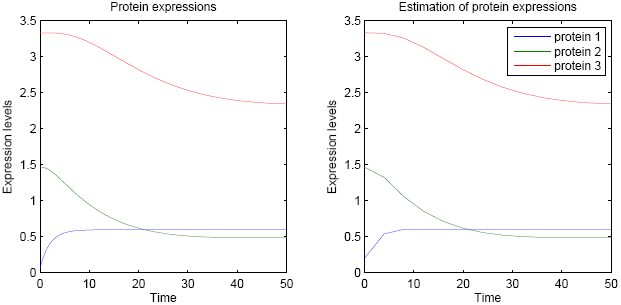
Exact protein expression curves derived from model ODEs (17) and (18) (left), and their recovered estimations using 12 unnormalized timepoint samples via (P2) (right). For convenience of graphical comparison, the values of *r*_*i*_ were drawn from the system equations. Protein expression is otherwise normalized with respect to *r*_*i*_, but this would result in a transformed scale for this qualitative comparison.

We finally examine Algorithm 1, (P3), for the goal of network recovery. In this scenario, our target is to estimate vectors **a**_*i*_ and **b**_*i*_. We assume that the degradation rates are known in advance and therefore, since the system does not contain any miRNA in this particular example, **λ**_*i*_ is completely at hand. Let us consider gene *3* where the true value of **a**_3_ = (0.1,0, 0.1,0,0,0,0) and **b**_3_ = (1,0,0.1, 0.1,0,0,0). Vectors **a**_3_ and **b**_3_ are indexed with regard to

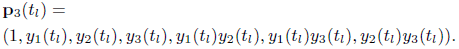

Applying our method, we obtain **a**_3_ ≃ (0.1, 0, 0.083,0,0, 0,0) and **a**_3_ ≃ (1,0,0.083, 0.08,0,0,0). Table I demonstrates that as the sampling frequency increases, we attain more accurate approximations. Furthermore, it can be seen that the estimations achieve similar accuracy after a small number of samples.

Employing the aforementioned single perturbation, we are only able to recover the strongest edge of gene 2, **b**_2_(6) = 10. The difficulty here is due to the sharp change in *y*1 (Figure 6), which provides us with a minimal amount of dynamic information. *y*1 near-instantaneously switches between two steady-state levels of expression, resulting in less accurate recovery of the underlying dynamics. However, expression patterns in perturbed biological settings tend to be more dynamic and are unlikely to contain this type of expression pattern. In this example, the removal of sharp instantaneous expression changes leads to complete recovery of the gene regulatory network.

**TABLE I.** Inference of binding coefficients describing energies of regulator complex-promoter interactions based on number of samples.

| # of Samples | Variables | Estimated vector entries |  |  |  |  |  |  |
| --- | --- | --- | --- | --- | --- | --- | --- | --- |
| 8 | $\mathbf{a}_3$ | 0.097 | 0 | 0.082 | 0 | 0 | 0 | 0 |
| | $\mathbf{b}_3$ | 1 | 0 | 0.082 | 0.06 | 0 | 0 | 0 |
| 16 | $\mathbf{a}_3$ | 0.1 | 0 | 0.093 | 0 | 0 | 0 | 0 |
| | $\mathbf{b}_3$ | 1 | 0 | 0.093 | 0.089 | 0 | 0 | 0 |
| 24 | $\mathbf{a}_3$ | 0.1 | 0 | 0.1 | 0 | 0 | 0 | 0 |
| | $\mathbf{b}_3$ | 1 | 0 | 0.1 | 0.092 | 0 | 0 | 0 |

System identification for gene 3 based on sample frequency. Coefficients $\mathbf{a}_3$ and $\mathbf{b}_3$ are the numerator and denominator of $p_3^{\text{bound}}$ (binding probability of RNA polymerase to a given promoter).

#### Remark 3.

*The recovery of regulatory networks using this proposed approach is tightly associated with the presence of dynamic changes in gene expression. These changes can provide us with a certain amount of information which predominantly specifies the accuracy of estimation. The achievable accuracy depends on many factors such as nonlinearity in changes or similarity in the range of changes.*

### B. Medium (10-gene) simulated network with noise

We extend our approach to simulated networks of 10 genes, generated as part of the DREAM4 *in silico* network inference challenge [42]. Each network dataset includes a simulated time series of gene expression in response to five chemical perturbations, along with single steady-state expression levels for wild-type, knockdown, knockout, and multifactorial perturbations. These datasets also simulate internal network noise and incorporate measurement noise. We use these data to assess the robustness of our approach in a non-ideal setup.

Our approach is geared towards precise genetic and chemical perturbations, while these datasets simulate chemicals that are nonspecific in their interactions. To place us at further disadvantage, we attempt network recovery using only the time series perturbations, forgoing all other datasets available to solvers. Lastly, our approach works best under conditions where RNA and protein degradation rates are known. Given that this information is unavailable, this exercise also serves as a test of our simplifying assumptions for such situations. Unlike simulations in the previous section, the rules of this challenge stipulate no self-regulation and no combined regulators.

DREAM4 Challenge 2 datasets for Networks 1 and 2 are used to infer gene regulatory networks and to inspect predictions of network topology using the official scoring pipeline. First, we use (P1) to produce smooth and continuous gene expression trajectories from the discrete and noisy time series datasets (Figure 7). Perturbed genes were identified and incorporated as described in Section III-B. Network inference is carried out using Algorithm 1. In the absence of RNA degradation rates, *λ_min_* is set to either 0.001 or 0.01, and *λ_max_* is set to 0.1 or 1. If a directed network edge is identified, the probability of the edge is set to 1 for weighted edges, and 0 otherwise. This is done to allow scoring of our network with the provided scripts, given our non-probabilistic formulation. Algorithm 1 minimization values are filtered against abnormal values that could represent underfitting and overfitting of data.

**Fig. 7.**
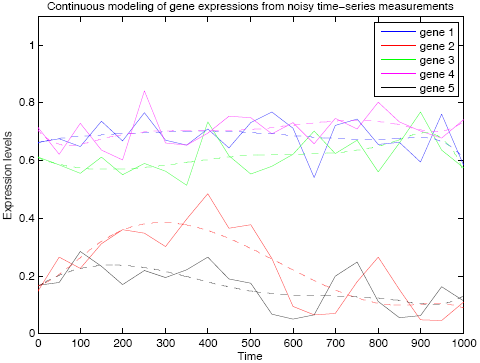
Time series gene expression measurements from simulated DREAM4 datasets are shown with connected solid lines. Dashed lines of corresponding color show that application of (P1) effectively produces noise-free (smooth) and continuous gene expression curves.

For Network 1, we report the area under the receiver operating characteristic curve (AUROC) = 0.81 and the area under the precision-recall curve (AUPR) = 0.75, and for Network 2, AUROC = 0.76 and AUPR = 0.68 (Supplemental Figure 1). These results compare very favorably to other time series-based methods applied to the same datasets [43]. In fact, for Networks 1 and 2, the AUROC and AUPR values represent improvements over the top reported results.

### C. Network inference from yeast cell cycle time series

In order to probe real biological data with inherent noise, we apply parts of our pipeline to a classical yeast cell cycle microarray dataset [44]. This data is provided as a 25 point time-series with a 5 minute sampling interval. Given the yeast cell is in an incredibly dynamic stage post synchronization with α-factor pheromone, this again represents a vast departure from ideal near steady-state conditions with a precise and local perturbation. We chose to focus our analysis on a set of primary regulatory genes and complexes involved in core cell cycle control and that showed greater than 15% changes in expression over the time course [41]. This led to retainment of 7 genes. We use (P2) to fit smooth continuous functions to the noisy gene expression measurements exhibited in Figure 8. We next examine our proposed scheme, (P4), to infer a gene regulatory network among these genes.

**Fig. 8.**
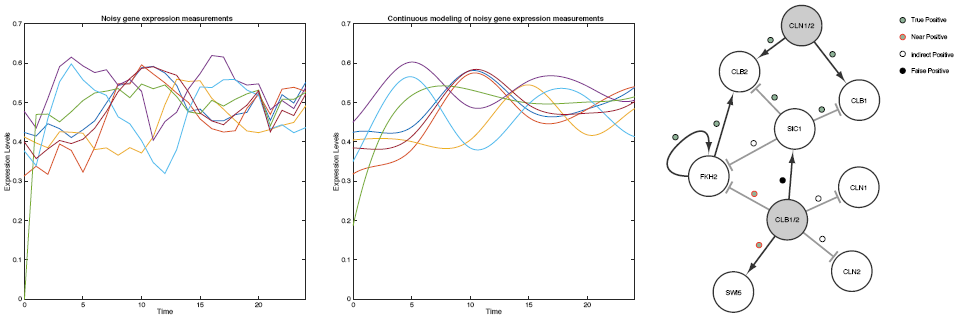
Time-series gene expression measurements of yeast cell cycle-associated genes filtered at a stringent change detection threshold ( T = 0:15) (left), and their recovered estimations using (P2) (center). The inferred network via (P4) is shown on the right, compared to the network as it’s presently understood ([41]). “True positives” represent edges recapitulated by the inference algorithm in both direction and influence, “near positives” represent edges correct in direction but with reversed influence, “indirect positives” represent edges of correct direction and influence with a missing intermediate node, and “false positive” indicates an edge not found in the reference network and that cannot be explained through a single intermediate node.

The inferred network is shown in Figure 8, with arrows indicating directed edges for gene-gene excitatory and inhibitory interactions. Of the 12 regulatory interactions inferred, 6 are correct in both directionality and influence (i.e. inhibition vs activation) and 2 are correct only in directionality. Further, 3 can be considered conditionally correct, whereby the predicted influence is mediated by a single intermediate node that was absent from the model. A single edge was labeled as a false positive, even though an argument can be made for mediation of that influence by two intermediate nodes. Strikingly, the algorithm correctly predicts a role for combined regulators and recovers the only example of self-regulation in the reference pathway. This is promising, given the absence of data relating to protein degradation, contextless inference, and the non step-wise nature of changes in expression that would be preferred in our proposed experimental scheme.

## V. Conclusions

The gene inference pipeline described in this work helps establish a robust framework for network discovery from perturbed expression data. The system of equations used to model eukaryotic gene regulation include the novel extension of a thermodynamic and statistical mechanic approach to polymerase binding. This pipeline is best suited for the processing of expression measurements from high-resolution time series experiments involving precise genetic or chemical perturbation of a steady state system. Genetic perturbation is best in the form of induced over-expression or RNAi-mediated gene knockdown. Chemical perturbation is best in the form of a chemical that has a specific protein interaction and limited off-target effects. However, we establish that this approach can yield insights under non-ideal conditions.

The modular nature of our pipeline allows for the modification of different stages to best fit a given biological system and of expression information. Alternative approaches can be implemented for the stags that precede the core inference algorithm, including change detection. The performance of this approach can further be improved with *a priori* knowledge of protein expression levels, protein and RNA degradation rates, along with the labeling of noncoding RNAs. Technologies are continually being improved for the purpose of capturing these data in a genome-wide manner [45], [46], [47], [48], to complement gene expression measurements. Our gene inference approach can readily utilize protein expression data, protein and RNA degradation data, and miRNA labeling data.

While we expect such inference approaches to work better for homogenous and synchronized single-cell or single-tissue systems, we also expect to capture the most prominent and meaningful aspects of the aggregate dynamics of heterogenous mixed-cell populations, multi-tissue systems, and whole organisms. Future directions include the more comprehensive validation and refinement of these algorithms for synthetic networks and higher-order eukaryotic systems, adaptations of more sophisticated change detection schemes, and surveys of a broader range of system-specific sampling frequencies.

This inference method has broad application in biological network discovery. For example, it can be used to identify the topology of gene regulatory networks immediate to drug response, and can be used to identify new interactions for genes implicated in disease. The inference data can then be used to seed and prioritize candidates for downstream biological and *in vivo* validation.

## Appendix

### A. Treatment of protein regulators

Consider a gene for which the probability of RNAP being bound to a specific promoter site, *S*, is under the potential influence of a single non-steady state regulator, Regulator 1, and the collection of all available regulators still in steady state. The steady state regulators are encapsulated as a single super-protein complex, *SS*, that is fixed as bound to the promoter region. Suppose that we have *P* RNAP, *R*_*1*_ Regulator 1, and *R*_*ss*_ super-protein complex.

We apply the following notation: 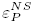 is used to denote the energy of the case in which RNAP is bound to a non-specific (*NS*) DNA binding site, 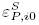 the energy when RNAP is only bound to the *S* binding site, 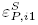 the energy when RNAP is specifically bound to the promoter-regulator complex, 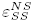 the energy when the *SS* is bound to the *NS* binding site, 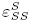 the energy when the *SS* is bound to the *S* binding site, 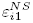 the energy when Regulator 1 is bound to the *NS* binding site, 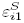 the energy when Regulator 1 is bound to the *S* binding site, and

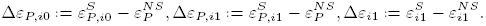

Also define

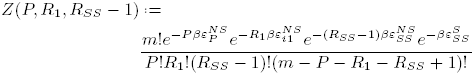

where *Z(P, R_1_, R_SS_ —* 1) gives the total number of arrangements for RNAP and R1 at *NS* binding sites, weighted by a Boltzmann factor providing a relative energy for each state.

The available configurations of the system with corresponding unnormalized probabilities are enumerated as follows: (i) Regulator *1* and RNAP unbound: *Z*(*P*, *R*_1_, *R*_*SS*_ — 1), (ii) only Regulator 1 bound: 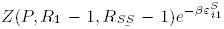, (iii) only RNAP bound: 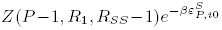, and (iv) both Regulator *1* and RNAP bound: 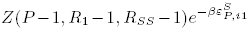. To derive the probability of RNAP binding, we sum the probabilities of configurations in which RNAP is bound to the specific site and divide over the sum of probabilities of all potential configurations, *Z*total. Here, in parallel to [22], it is shown how the effect of steady state proteins can effectively be removed from the protein regulator formulation, under the aforementioned arrangement. To represent the probability of RNAP binding to the cis regulatory region of gene *i*, we define 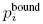 as follows.

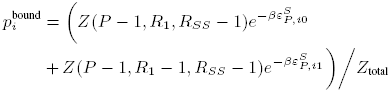

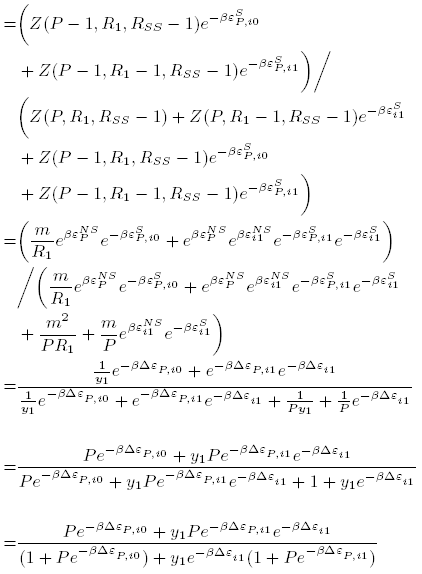

where we have applied the approximation *m*!/*P!R*_1_!(*R*_*SS*_−1)!(*m*− *P* − *R*_1_ − *R*_*SS*_ + 2)! ≈ *m*^*P*^ *m*^*R*1^*m*^*RSS*^/*P*!*R*_1_!(*R*_*SS*_ − 1)!. We introduce *y1*, the protein product of Regulator 1 defined as *R*_1_/*m*, for the purposes of normalization and in keeping with the protein designations used throughout this paper. We additionally note that *P* in the final steps of the derivation above is also normalized to *m*, but we retain the same notation for simplicity.

The final derivation can be generalized to, for an indefinite number of first and second order regulators.

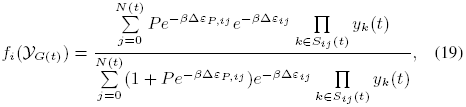

where Δ*ε_ij_* is the binding energy of the *j*th complex to the promoter, Δ*ε_P,ij_* is the energy of RNAP being bound to the promoter-regulator complex *j*, and *P* is the concentration of RNAP. Setting *a*_*ij*_ = *P*_*e*_^−βΔε*P*,*ij*^e^−βΔε*ij*^ and *b*_*ij*_ = (1 + *Pe*^−βΔε*P*,ij^)*e*^−βΔε*ij*^, we arrive at the form given in (3).

### B. B-splines

B-splines have been well investigated in approximation theory and numerical analysis, leading to a variety of important properties such as computational efficiency and numerical stability. Particularly, the B-spline basis functions have the best approximation capacity based on the Stone-Weierstrass Approximation Theorem. Polynomial functions are also used to estimate continuous functions. However, the B-spline bases are shown to be optimally stable [49].

A set of B-spline basis functions in variable *t* is determined by the degree of a piecewise polynomial, *P*, and a knot sequence [50]. The knot sequence is a set of points that divides a real interval into a number of sub-intervals. More precisely, *D* bases of degree *P* are parameterized by *D* + *P* + 1 knots, {*t*_0_, *t*_1_,…, *t*_D+P_} where *t*_0_ ≤ *t*_1_ … ≤ *t*_*D+P*_. Employing this set of knots and the De Boor recursion in [51], the *d*th B-spline basis of degree *P*, written as 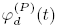, is derived recursively as follows:

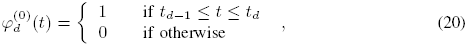

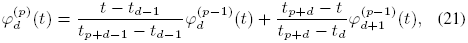

for 1 ≤ *d* ≤ *D + P* — *p* where *p* = 0 in (20) and 1 ≤ *p* ≤ *P* in (21). The above recursion is visualized in Figure 9 (reconstructed from [50]).

**Fig. 9.**
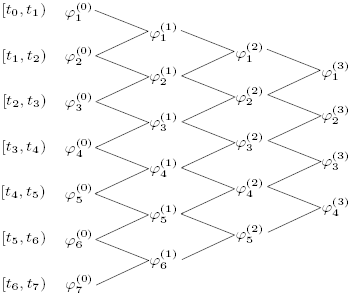
The De Boor recursion for *P* = 3 and *D* = 4.

The degree *P* = 3 or *4* is sufficient in most applications. The number of basis functions should be large enough to arrive at accurate estimation but not too large to cause overfitting. In our case, gene and protein levels do not contain high frequency changes and therefore, a small number of basis functions are sufficient to represent gene and protein expressions.

### C. Bi-Convex Problems

Bi-convex optimization is a generalization of convex optimization where the objective function and the constraint set can be bi-convex [39].

#### Definition 1

*Let x* ⊆ ℝ^n^ *and y* ⊆ ℝ^n^ *be two non-empty convex sets. The set E ⊆ χ × γ is called bi-convex if B_x_* := {*y ε γ* : (*x,y*) ε *B*} *is convex for each *x*, and *B*_*y*_ := {*x ε χ* : (*x,y*) γ B*} *is convex for each y*.

#### Definition 2

*A function f*(*x, y*) : *B* → ℝ *is called bi-convex if f* (*x, y*) *is convex on B_x_ for every fixed x and also convex on B_y_ for every fixed y.*

A common method to solve a bi-convex problem is ADMM [52]. The ADMM is an iterative augmented Lagrangian method that uses partial updates for dual variables and replaces joint minimization by simpler sub-problems. However, the mentioned procedure does not guarantee global optimality of the solution.

### D. Proof of Theorem 1

The stationary points 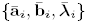 of (P3) are derived by setting sub-gradients to zero as follows

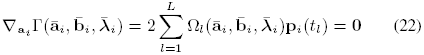

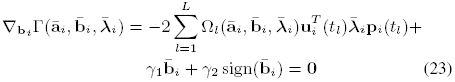

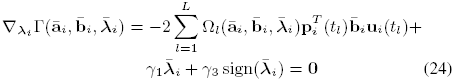

with respect to constraints 0 ≤ **α**_*i*_ ≤ 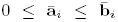 ≥ 0. These constraints admit that sign(·) can be replaced by vector 1 in the above equations. It is obvious from (23)-(24) that 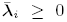, which results in 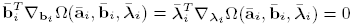, which results in

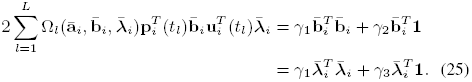

Consider the convex optimization

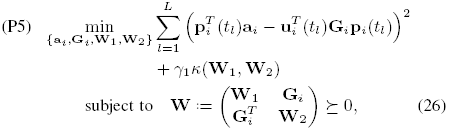

where 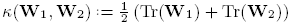.

Minimizing (P5) with respect to {**W**_1_, **W**_2_} leads to

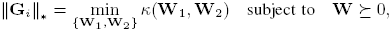

which is the alternative characterization of the nuclear norm [53]. Taking advantage of the nuclear norm, we can restrict matrix G_i_ to be rank one as 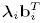. Also, *κ(·, ·)* is able to satisfy the required sparsity for 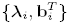. To investigate these claims, recall constraints (??) and set 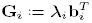, and 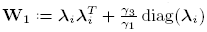, and 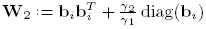 where diag(λ_i_) is the diagonal matrix with *(j, j*)th entry equal to *λ_i_(j).* Then, the triple (G_i_, W_1_, W_2_) is feasible for (P5) due to

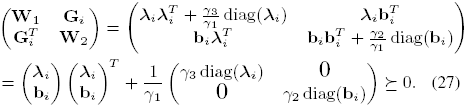

In addition, we have

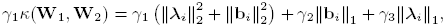

and therefore the same objective function for (P3) and (P5) are obtained. This proves any feasible solution of (P5) yields an inner bound for (P3).

We next establish that the proposed inner bound is always equal to (P3) upon satisfying the condition introduced in Theorem 1 and conclude the two problems are equivalent. The equivalence ensures that the stationary point of (P3) (which exhibits Theorem 1 condition) is in fact globally optimal. To show this, the Lagrangian is first formed as

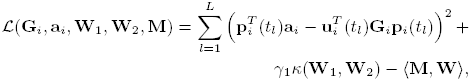

and M indicates the dual variable associated with the constraint **W** ? 0 tn accordance with the block structure of **W** in (P5), we define *Mi* = *[M*]_11_, *M2 .= [M]_12_, M3* = *[M]_22_,* and *M4 = [M]_21_*. The optimal solution of (P5) must

i. null the sub-gradients

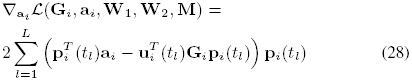

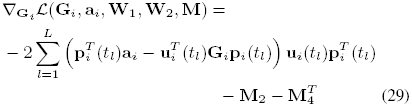

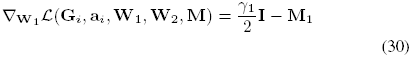

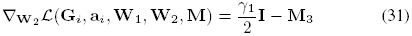

and also satisfy

i. the complementary slackness condition (M, W) = 0;
ii. primal feasibility **W***≥* 0;
iii. dual feasibility M ≥ 0.

Consider the stationary points of (P3), and choose the candidate primal variables 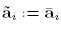 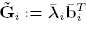, 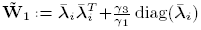, 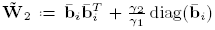; and the dual variables 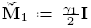, 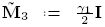, 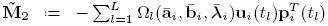 and 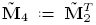. Then, condition (i) holds because the sub-gradients (28)-(31) are zero when substituting the introduced primal and dual variables. The requirement (ii) is also true since

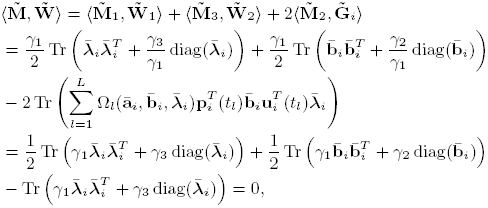

where the last equality follows from (25). Moreover, (iii) is confirmed similar to (27). In order to meet the last criterion (iv), according to a Schur complement argument [37], it is sufficient to invoke 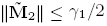.

Consequently, by choosing the proposed candidates that have been proved to be optimal, one can easily verify (P5) coincides with (P3). This completes the proof.

---

**supplementary Figure 1.**
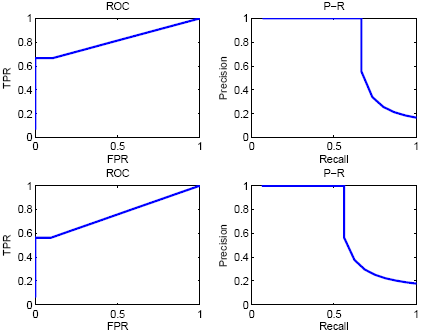
RoC and P-R curves for Dream 4, Challenge 2 Network 1 (top) and Network 2 (bottom).

